# High school science fair: Positive and negative outcomes

**DOI:** 10.1101/737270

**Authors:** Frederick Grinnell, Simon Dalley, Joan Reisch

## Abstract

The goal of our ongoing research is to identify strengths and weaknesses of high school level science fair and improvements that can help science educators make science fair a more effective, inclusive and equitable learning experience. In this paper, we confirm and extend our previous findings in several important ways. We added new questions to our anonymous and voluntary surveys to learn the extent to which students had an interest in science or engineering careers and if science fair participation increased their interest in science or engineering. And we surveyed a national rather than regional high school student group by incorporating our survey into the Scienteer online portal now used by Texas and some other states for science fair registration, parental consent, and project management. We learned that about ~60% of the more than 300 students in the national cohorts completing surveys in 2017 and 2018 said that they were interested in a career in science or engineering, and ~60% said that participating in science fair increased their interest in science or engineering. About two-thirds of the students were required to participate in science fair, and that requirement reduced the frequency of students who said that science fair increased their interest. In the worst case, ~10% of the students who said that they were not interested in a career in science or engineering and required to participate in science fair engaged in research misconduct (i.e., plagiarism and making up their results). Students’ positive comments about competition in science fair focused on the competition incentive, whereas their positive comments about science fair that was non-competitive focused on learning about the scientific process and learning in general. We discuss the findings in the context of National Science Teaching Association guidance about voluntary science fair participation and begin to identify features of science fair practice consistent with increased student interest in the sciences or engineering.

## Introduction

Next Generation Science Standards (NGSS) identifies experiencing the practices of science as one of three essential dimensions of science education, “students cannot comprehend scientific practices, nor fully appreciate the nature of scientific knowledge itself, without directly experiencing those practices for themselves” [1]. The question how to integrate the practice of science into science curricula is not new. Debates about how to do so permeate the history of science education [2]. Science fairs offer students an attractive opportunity to experience the practices of science for themselves because students who participate go through the processes of selecting a problem and question to answer; designing and implementing experiments to answer the question; analyzing and drawing conclusions from the results; and explaining the findings to others through interviews and poster presentations [3–7].

Science fairs receive a lot of public attention. President Obama stated in his 2011 State of the Union Address, *We need to teach our kids that it’s not just the winner of the Super Bowl who deserves to be celebrated, but the winner of the science fair* [8]. The film *Science Fair* won the 2018 Sundance Film Festival *festival favorite award*. A 2019 GEICO television commercial “Science Fair of the Future” had more than 11 million views on YouTube in its first month. Nevertheless, despite the long history and wide implementation as part of informal and formal science education in the United States, few published research studies examine how science fair participation affects student engagement with science [7]. National Science Teaching Association (NSTA) guidance takes the position that student participation in science fairs should be voluntary with emphasis placed on the learning experience rather than on the competition [9]. However, whether most students who participate in high school science fair are required or choose to participate and to what extent the students perceive science fairs as emphasizing learning vs. competition are open research questions.

The overarching hypothesis guiding our research is that a better understanding of science fair practices will help science educators make science fair a more effective, inclusive and equitable learning experience. Rather than theoretical, our aim is to improve the practical implementation of science fairs based on an analysis of students’ high school science fair experiences. We began our research during 2014, conducting surveys with a group of regional high school students who had just competed in the Dallas Regional Science and Engineering Fair (DRSEF) and with post high school students on biomedical science educational trajectories doing research at UT Southwestern Medical Center. The post high school students may or may not have participated in science fair. The surveys were anonymous and voluntary and characterized student experiences by asking them in addition to demographic information to identify sources of help they received, types of help received, obstacles encountered, and ways of overcoming obstacles [10, 11].

In this paper, we confirm and extend our previous findings in several important ways. First, we added new survey questions to learn the extent to which students had an interest in science or engineering careers and if science fair participation increased their interest in science and engineering. Second, we surveyed a national group of high school students by incorporating our survey into the Scienteer (www.scienteer.com) online portal now used by Texas and some other states for science fair registration, parental consent, and project management. We found that about 2/3 of the students in the national cohort who completed surveys in 2017 and 2018 had been required to participate in science fair and observed negative consequences of requiring participation on student science fair experiences and attitudes. Some policy implications of the latest findings have been put forth in a NSTA Reports commentary [12].

## Materials and methods

This study was approved by the UT Southwestern Medical Center IRB (#STU 072014-076). Study design entailed administering to students a voluntary and anonymous online survey [10, 11] using the REDCap survey and data management tool [13]. Survey content, adapted from earlier research by others [14], was similar overall as our previous studies [10, 11] and included questions about student demographics, type of science fair participation, help expected and received, and obstacles encountered and solutions implemented to overcome obstacles. Also, the survey used in the current studies included new questions about student interest in a career in the sciences or engineering and the impact of science fair participation on interest in science. The survey can be found in supporting information (S1_Survey).

High school students were invited to participate in the science fair survey through the Scienteer (www.scienteer.com) online portal used in Alabama, Maine, Missouri, Texas, Vermont, and Virginia for student science fair registration, parental consent, and project management. After giving consent for their students to participate in science fair, parents could consent for their students to take part in the science fair survey. To prevent any misunderstanding by parents or students about a possible impact of agreeing to participate or actually participating in the survey, access was not available to students until after they finished <u>all</u> of their science fair activities. Students were instructed to log in to Scienteer after completing the final science fair activity in which they participated. Those who did so were presented with an alert and hyperlink to the science fair survey. No incentives were offered for participation, and Scienteer does not send out reminder emails.

Table 1 summarizes the student survey response rate. Of the students who clicked on the hyperlink, 20-25% completed the surveys. We don’t know if some students logged back into Scienteer but did not click on the hyperlink so the maximum response rate would have been ~20%. Overall, students who completed surveys represented slightly more than 0.5% of all students who signed up for science fair through Scienteer. Given that participation in the survey involved an indirect, single electronic invitation without incentive or follow-up, a low response rate was not surprising [15–17]. Most of the submitted surveys (>90%) were complete and non-duplicates. These surveys were used for data analysis. The complete survey data sets for students who participated during 2017 and 2018 school years can be found in supporting information (S1_Dataset and S2_Dataset).

**Table 1.**
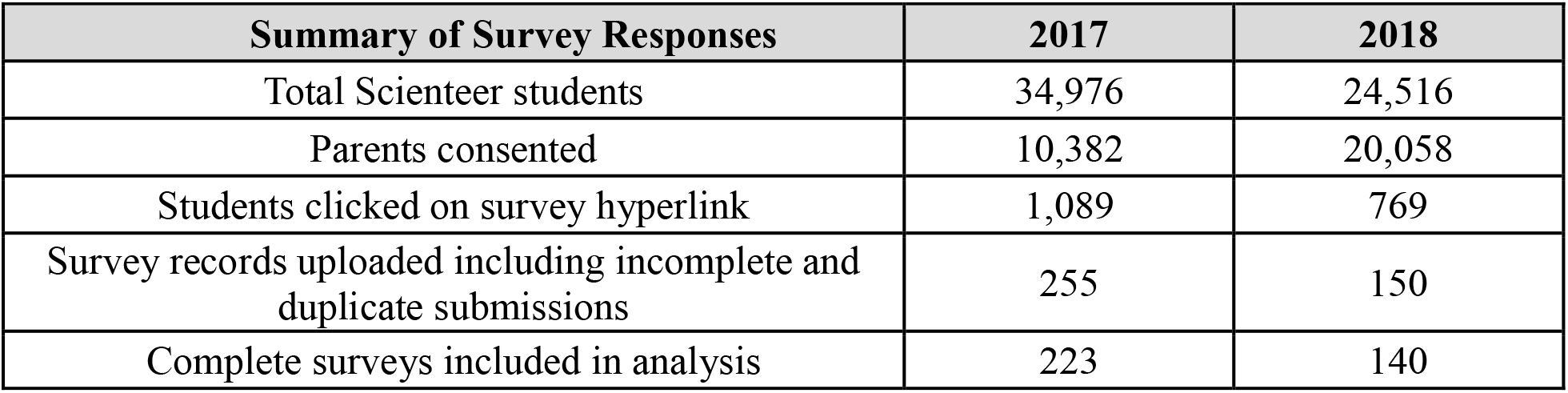
Student survey responses.

Quantitative data were analyzed by frequency counts and percentages. Data were sorted to compare different answer selections. Significance of potential relationships between data items was assessed using relevant statistical methods, e.g., Chi-square contingency tables for independent groups. Results shown in the figures are presented two ways -- graphically to make overall trends easier to appreciate and in tables beneath the graphs to show the actual numbers. A probability value of 0.05 or smaller was accepted as statistically significant but actual p values are indicated where significant differences were observed. Results for 2017 and 2018 national cohorts are shown separately in Figs 1-3 and S1-4 Figs but otherwise combined.

**Figure 1.**
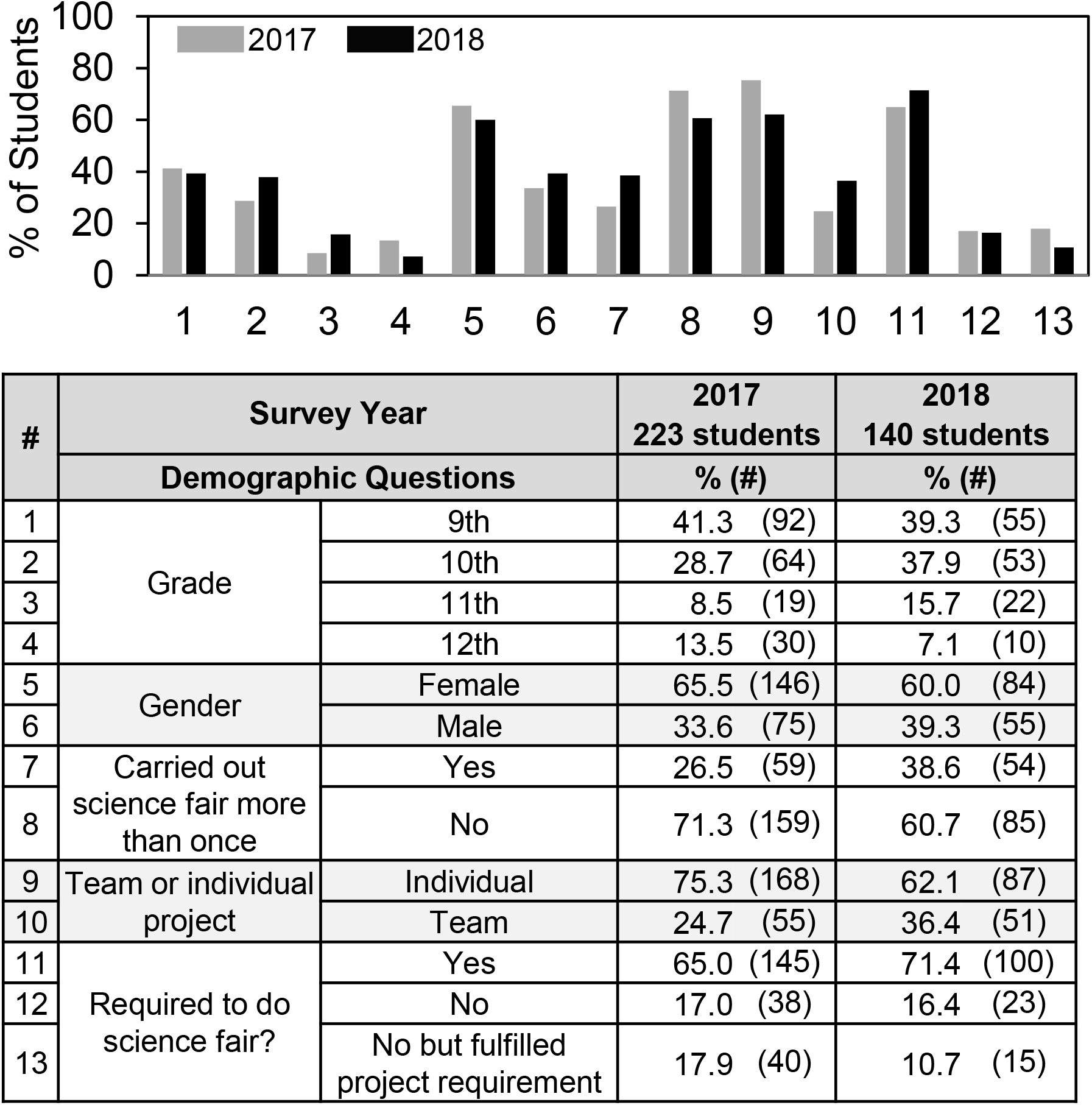
Student demographics. Summary of student survey demographic information. Most but not all students answered every demographic question.

Qualitative text analysis for the open-ended text questions was accomplished as described previously [11] using an approach modeled on NVivo [18, 19] based on grounded theory [20]. More than 80% of the students who completed surveys wrote comments about why science fairs should be optional or required. Two members of the research team (FG and SD) independently coded students’ comments, which were categorized into a matrix of shared student reasons (nodes). The independently coded matrices were revised and harmonized into 16 *Reason Why* categories why science fair should be required or optional. Longer student comments frequently expressed more than one idea, in which case the comments were coded into more than one *Reason Why* category, and which is why the number of reasons exceeds the total number of student comments. The complete set of student answers to the *Reason Why* question and corresponding reason category assignments can be found in supporting information (S3_Dataset).

## Results

### Survey demographics

Fig 1 shows the similarity of student responder cohorts in 2017 and 2018. Most students who participated in the survey (~75%) were in 9^th^ and 10^th^ grades. More girls than boys completed surveys. About one in three students had carried out science fair more than once. The surveys are anonymous; therefore, we do not know if any students who completed surveys in 2017 also did so in 2018, but the survey instructs the students that if they carried out science fair more than once, then they should answer the survey questions according to their most recent experiences. Three out of four student projects were individual.

Overall, 65-70% of the students who participated in science fair reported that they were required to do so. Since the survey does not provide ancillary information regarding what it means for science fair to be required, the students’ answers reflect how they felt about their participation. We cannot tell if they understand “required to do science fair” differently from their schools’ intentions, e.g., required to participate in science fair to get into an advanced class or to increase one’s grade is not the same as truly required but can be perceived that way.

### Student experiences in high school science fair–help and obstacles

Student answers to questions regarding sources of help, types of help received, obstacles encountered, and ways of overcoming obstacles were very similar comparing the 2017 and 2018 responders. Fig 2 presents a graphical summary of the results with details in the corresponding supplemental figures S1-S4 Figs.

**Fig 2:**
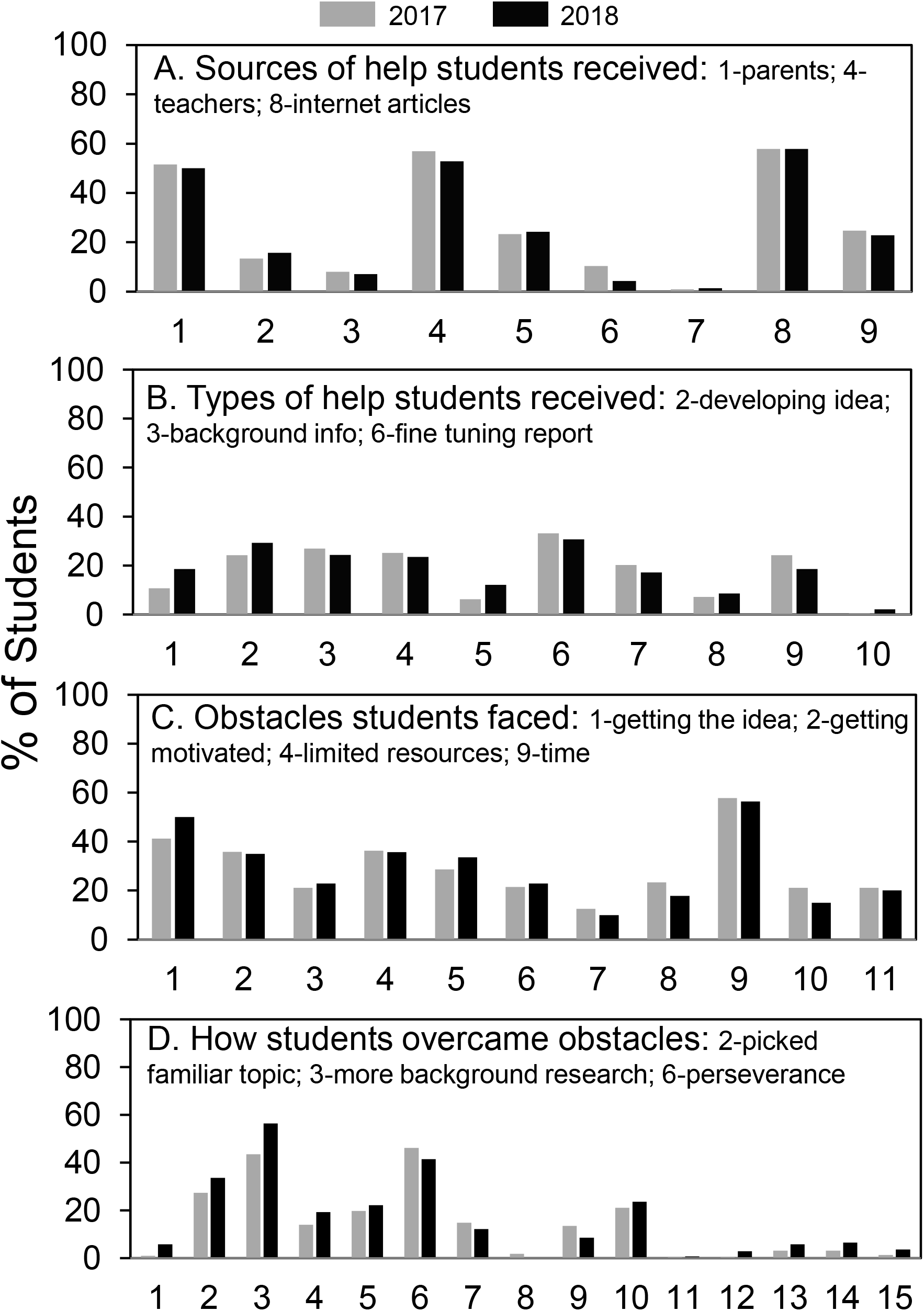
Summary of student science fair experiences regarding help and obstacles. Data are summarized from S1-S4 Figs.

The most frequent student selections are labeled. (A) Parents, teachers, and articles on the internet were the main sources of help reported by more than 50% of the students. (B) No more than 35% of the students reported receiving any particular type of help with the most frequent types of help received developing the idea, background information and fine-tuning the report. Even though only about a third of students received any particular type of help, a large majority of students reported receiving the kind and amount of help that they wanted from teachers (see S2 Fig). (C) Regarding obstacles faced, the most frequent selections were getting the idea, getting motivated, limited resources, and (above all) time. (D) Overcoming obstacles was accomplished most often by picking a familiar topic, doing more background research, and perseverance. Five of the students indicated that they used someone else’s data (D, #12) and 15 said they made up their data (D, #13) (see S4 Fig).

### Comparison of National and Regional Student Experiences

In Table 2, we compare the most frequent selections by the 2017-2018 national student groups (averaged) with data previously published based on surveys of regional students [10]. Most of the top choices (item rank) of the national and regional groups overlapped in every category. One major difference was that 85% of the regional students reported receiving coaching for the interview compared to only 21% of the national students. This difference and several others -- use of articles in books and magazines, more background research, and more perseverance -- are consistent with the highly supportive practices by the North Texas suburban school district where most of the regional students attended high school, and where students were incentivized rather than required to participate in science fair. Indeed, only 8% of the regional students were required to participate in science fair compared to 68% of the national students.

**Table 2.** Comparison of highest ranked selections from national and regional student survey results.

| Survey Item |  | Student Group |  |  |  |
| --- | --- | --- | --- | --- | --- |
|  |  | National<br>% (#) of 363<br>students | Item<br>rank | Regional<br>% (#) of 64<br>students | Item<br>rank |
| Sources of help<br>received<br>(9 options) | Articles on internet | 57.9 (210) | 1 | 53.8 (35) | 3 |
|  | Teachers | 55.4 (201) | 2 | 69.2 (45) | 1 |
|  | Parents | 51.0 (185) | 3 | 41.5 (27) | 4 |
|  | Articles in Books or Magazines | 24.0 (87) | 4 | 64.6 (42) | 2 |
| Types of help<br>received<br>(10 options) | Fine-tuning report | 32.2 (117) | 1 | 47.7 (31) | 2 |
|  | Developing idea | 26.2 (95) | 2 | 29.2 (19) | 4 |
|  | Background information &<br>finding research site and<br>participants | 25.9 (94) | 3 | 36.9 (24) | 3 |
|  | Coaching for the Interview | 22.0 (80) | 5 | 84.6 (55) | 1 |
| What obstacles did<br>you encounter?<br>(11 options) | Time pressure | 44.6 (162) | 1 | 64.6 (42) | 1 |
|  | Coming up with the idea | 57.3 (208) | 2 | 63.1 (41) | 2 |
| How did you<br>overcome<br>obstacles?<br>(15 options) | More background research | 48.5 (176) | 1 | 72.3 (47) | 1 |
|  | Perseverance and self-discipline | 44.4 (161) | 2 | 69.2 (45) | 2 |
| Required to do science fair |  | 67.5 (245) |  | 7.7 (5) |  |

### Effect of Science Fair on Student Interest in a Career in the Sciences or Engineering and the Consequences of Requiring Science Fair Participation

An important positive outcome of science fair would be for students to become more interested in science. Fig 3 presents an overview of student answers to two related questions, one regarding the students’ interests in a career in the sciences or engineering, and the other regarding whether science fair participation increased their interest in science or engineering. About 60% of the students overall said they were interested in a career in the sciences or engineering; 15% said they were not; and the remainder were unsure. Also, about 60% of the students said that science fair participation increased their interest in the sciences or engineering.

**Fig 3:**
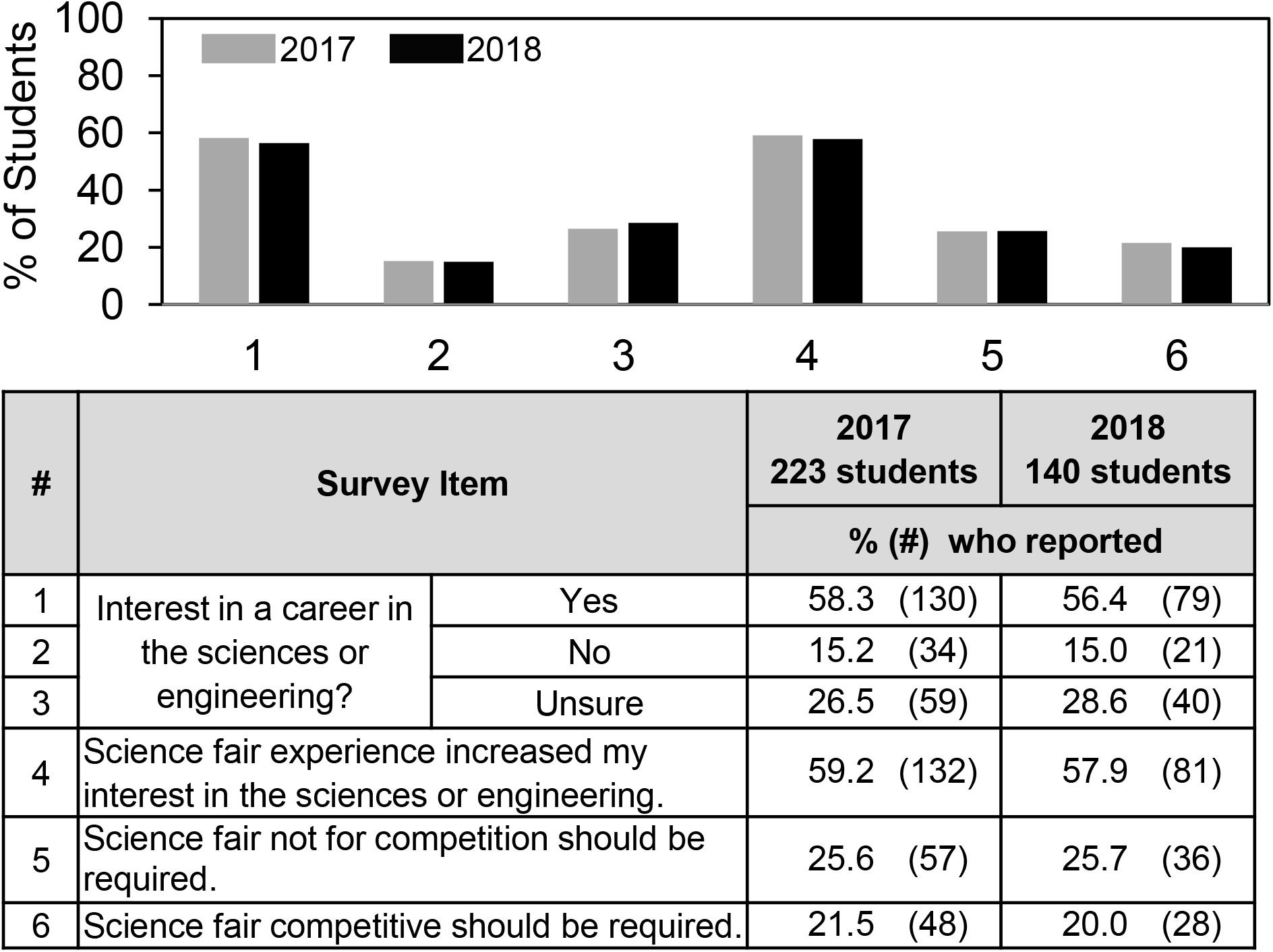
Frequency of student answers to the questions regarding student interest in a career in the sciences or engineering, impact of science fair on interest in science, and attitude towards requiring science fair.

As an indirect means to assess how students viewed the value of science fair, we asked the quantitative question: *Do you think science fair should be required or optional?* and the qualitative, open-ended text question: *Reason Why?* And we asked these questions for both competitive and non-competitive science fair to provide insights about student attitudes towards competition *per se*. Fig 6 shows the quantitative finding. Similar to previously reported results for the regional high school students [11], only 1 in 5 of the national students favored requiring science fair competition. That number was marginally but not significantly higher if science fair was described as non-competitive vs. competitive. Qualitative results of the open-ended text question will be described later.

Fig 4 shows some differences that reached significance comparing students who said that science fair did vs. did not increase their interest in science. Not surprisingly, the impact of science fair participation on student interest paralleled student attitudes towards a career in the sciences or engineering. In addition, students who reported that science fair increased their interest in science or engineering were more likely to have received help from teachers, from articles in books and magazines, and coaching for the interview. More of these students did additional background research and they reported more perseverance and self-discipline. Conversely, students required to do science fair were less likely to say that science fair participation increased their interest in the sciences and engineering and more likely to report that getting motivated was an obstacle

**Fig 4:**
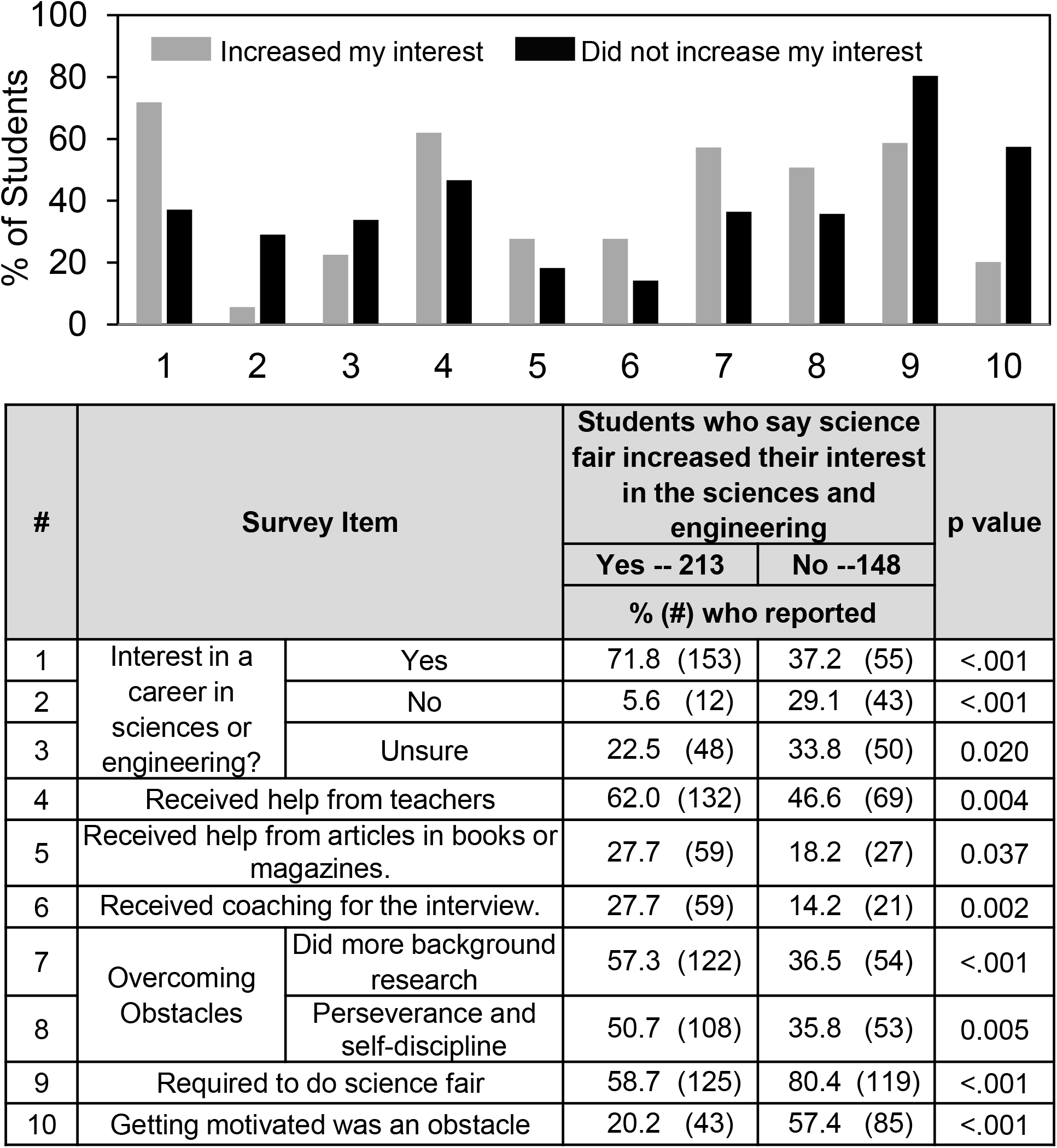
Differences in student experiences depending on whether students said that science fair increased their interest in science.

Figs 5 and 6 show more clearly the negative impact of requiring science fair. Fig 5 shows that regardless whether or not students were interested in a career in science or engineering, requiring them to participate in science fair decreased the number who said that participating in science fair increased their interest. Fig 6 shows that students who were required to participate in science fair were more likely to use someone else’s data or make up their data. Overall, ~10% of the students who also said they were not interested in a career in the sciences or engineering and were required to participate in science fair did one or the other. Rather than becoming more interested in science, these students committing research misconduct, i.e., using someone else’s data or making up their data. Doing so represents a serious negative outcome of participating in science fair.

**Fig 5:**
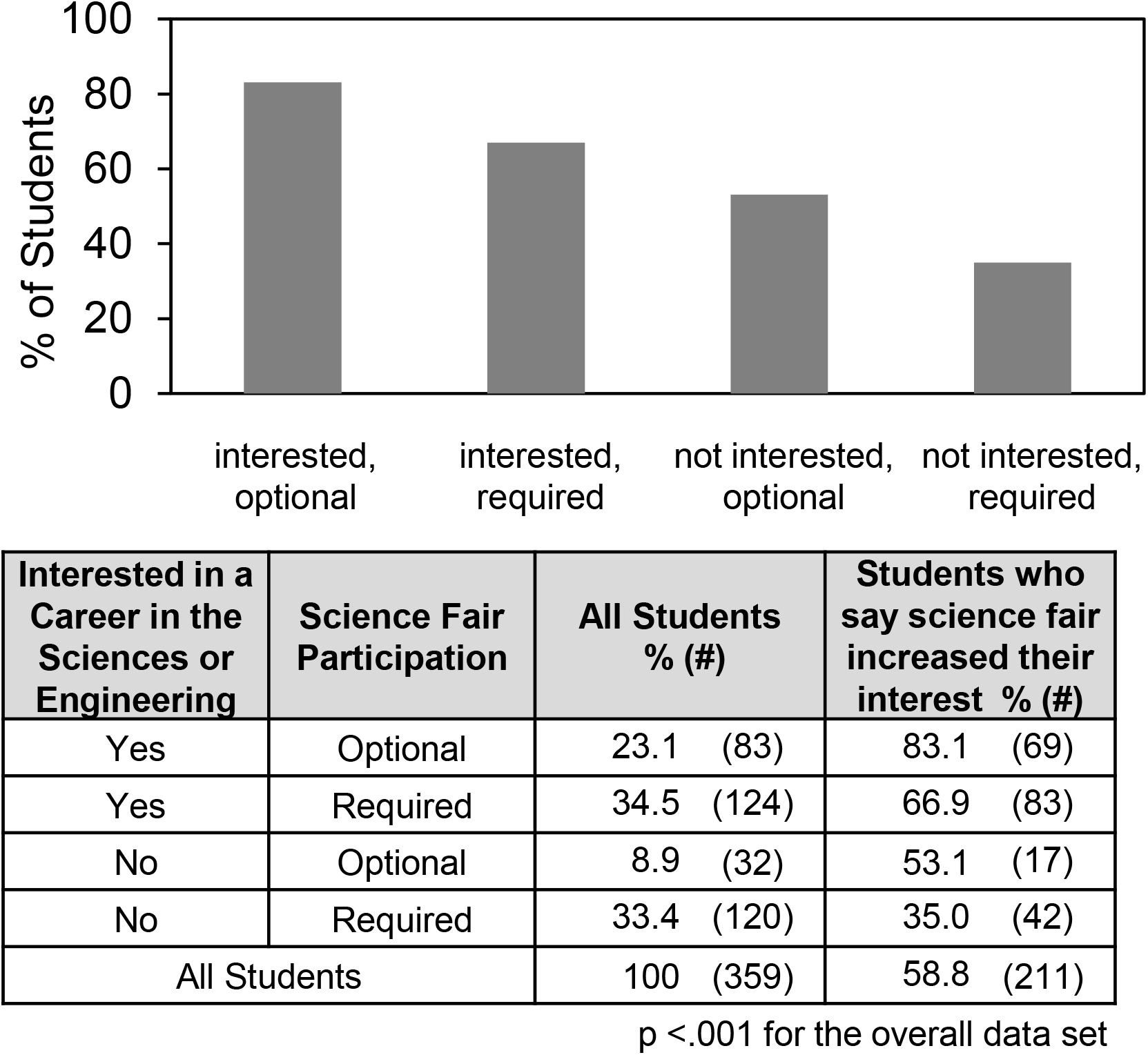
Students who say science fair increased their interest in science according to their interest in a career in the sciences or engineering and science fair requirement

**Fig 6:**
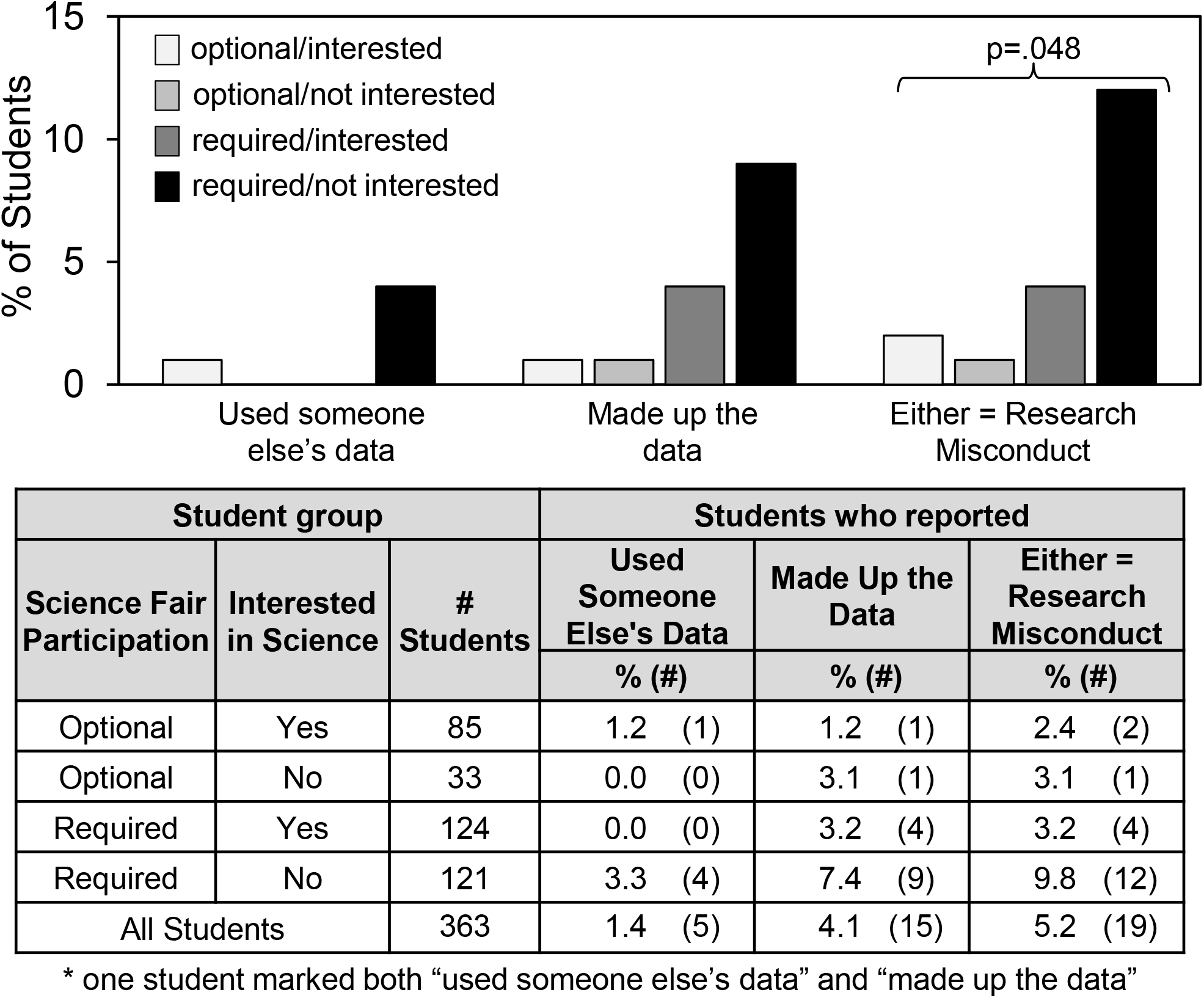
Research misconduct by students depending on science fair requirement and interest in a career in the sciences or engineering.

### Student Reasons -- Qualitative findings

Given the voluntary and anonymous format of our surveys, interviewing students was not a possibility. However, the open-ended text questions asking students to state reasons why science fair should be optional or required provided a rich source of insights regarding student attitudes. A total of 314 students (86.5%) commented about non-competitive science fair and 301 students (82.9%) commented about competitive science fair regarding why science fair should be optional or required. That more than 80% of the students wrote thoughtful answers was one indication that the students took the surveys seriously.

Two members of the research team (FG and SD) independently coded the comments, which were categorized into a matrix of shared student reasons (nodes). The independently coded matrices were revised and harmonized into 16 *Reason Why* categories that contained 445 student reasons about non-competitive science fair and 378 student reasons about competitive science fair. Table 3 shows the 16 *Reason Why* categories (7 positive and 9 negative) and examples of the students’ comments. Longer comments frequently expressed more than one idea, in which case the comments were coded into more than one *Reason Why* category. For instance, the student comment, *Science Fairs encourage students to learn new things in science in specific areas that interest them, which might lead to a future career in the science department*, was placed into both the “Introduction to scientific knowledge” and “Career interests” categories.

**Table 3.** Student reasons about science fair requirements organized according to positive (P) and negative (N) reasons with examples.

| P/N | Reason Category | Examples -- When a single comment is used to express more than one idea, the text relevant to the idea in bold italics. |
| --- | --- | --- |
| P | Intro to the scientific process | It helps enhance the student's knowledge of conducting an experiment, the scientific method, and a subject outside of school. |
| P | Communication or presentation skills | It builds up one's ability to present one's scientific findings or observations. |
| P | Intro to scientific knowledge | Science Fairs encourage students to ***learn new things in science in specific areas that interest them***, which might lead to a future career in the science department. |
| P | Career interests | Science Fairs encourage students to learn new things in science in specific areas that interest them, which ***might lead to a future career in the science department***. |
| P | Competition incentive | Competition is a motivation for many students who want to be known as the best. |
| P | General learning | It is a good opportunity for students to build their education level and thinking. |
| P | Other positive | Some people do not want to spend a lot of time working on a project that they don't want to do, but I myself had a lot of fun doing it. |
| N | Not everyone interested in science | Because not everyone has a passion for science |
| N | Too much stress/pressure | I feel that many kids are stressed with other assignments... one more project is the last thing they need |
| N | No enjoyment and negative attitude | Because people shouldn't be **forced to do something they don't want to do**. Also people might not have the time or resources to do a science project. |
| N | Negative behaviors and consequences | There are many people who put forth no effort in their projects and make up data. |
| N | No time/money | Because people shouldn't be forced to do something they don't want to do. Also people **might not have the time or resources** to do a science project. |
| N | No value | Doing a basic project, like most people do, is more harmful then helpful, and serves no purpose other than wasting time |
| N | Don't like to compete | Some people just like to research and not compete |
| N | Don't like to make public presentations | Not all students may be comfortable speaking in front of people. |
| N | Other negative | They're [sic] certain way of having things done isn't the way everybody normally works. |

Fig 7 shows the frequency with which the positive and negative reasons were mentioned. The order of reasons is the same as in Table 3. Negative reasons outnumbered positive ones for both non-competitive (314 vs. 131) and competitive (277 vs. 101) science fair, but the reason categories differed. For non-competitive science fair, the most frequently mentioned negative reasons were “No enjoyment/negative attitude” (~22% of the students) and “No time/money” (~17% of the students); whereas for competitive science fair, the most frequently mentioned negative reason was “Don’t like to compete” (~22% of the students). The most frequently mentioned positive reasons for non-competitive science fair were “Introduction to the scientific process” and “General learning” (each ~8% of the students) vs. “Competition incentive” (~14% of the students) for competitive science fair.

**Fig 7.**
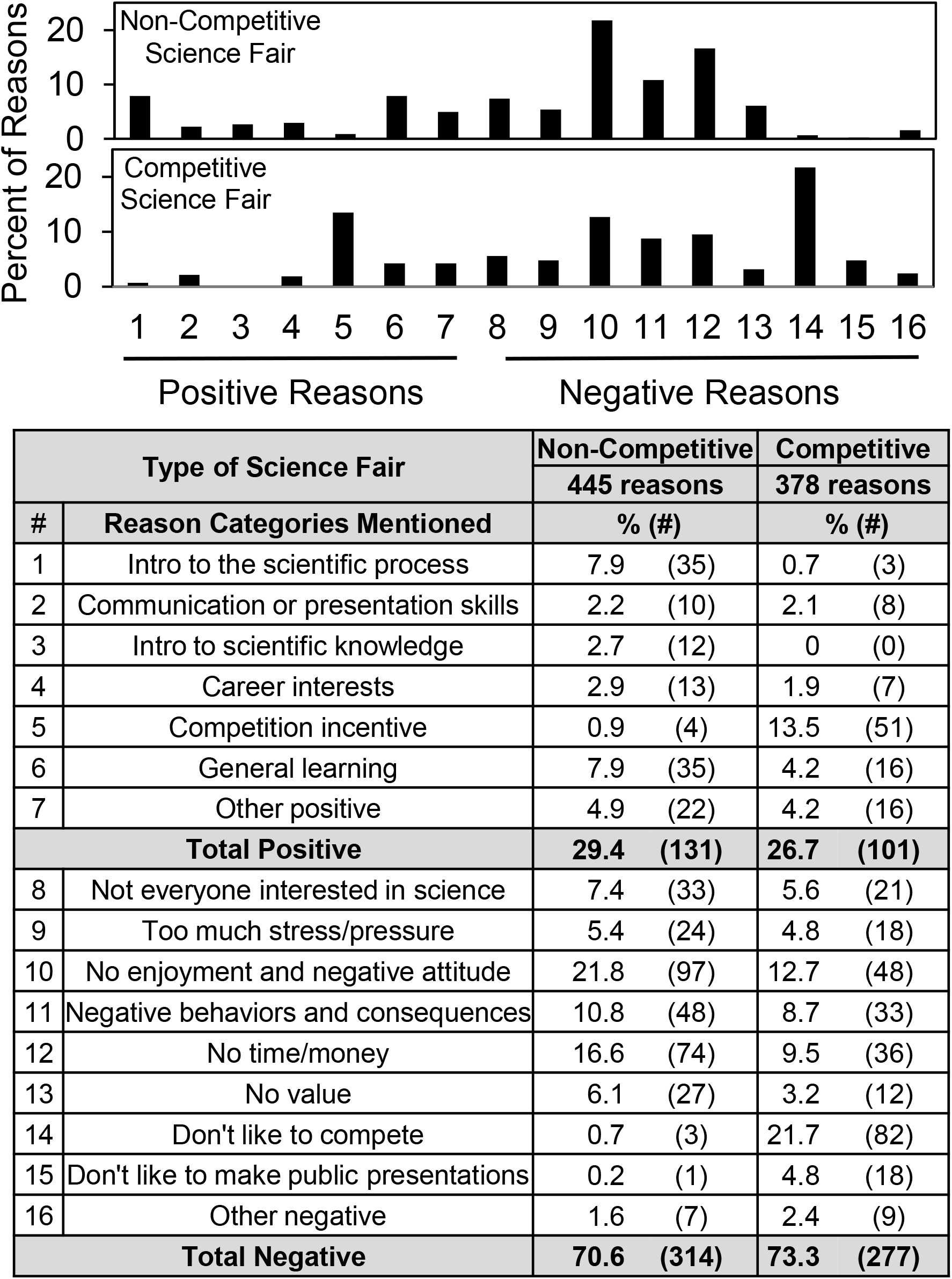
Distribution of student reasons positive and negative to require science fair.

If the results in Fig 7 were sorted according to students’ quantitative responses to the question whether or not science fair should be required, then 21-26% of students who said that science fair should be required accounted for 98% of the positive reasons regarding non-competitive science fair and 95% of the positive reasons regarding competitive science fair (S5 Figure). That the students’ open-ended comments compared favorably to their quantitative answers demonstrated internal survey consistency.

Fig 8 sorts the results in Fig 7 according to students’ quantitative responses to the question whether participating in science fair increased their interest in science. Students who said that science fair increased their interest in science or engineering were more likely to write positive comments in every category, especially introduction to process of science and general learning. Also, these students were more likely to select “competition incentive” for competitive science fair. On the other hand, students who said science fair increased their interests had more complaints about negative behaviors and consequences regarding non-competitive science fair and about disliking having to make a public presentation for competitive science fair.

**Fig 8:**
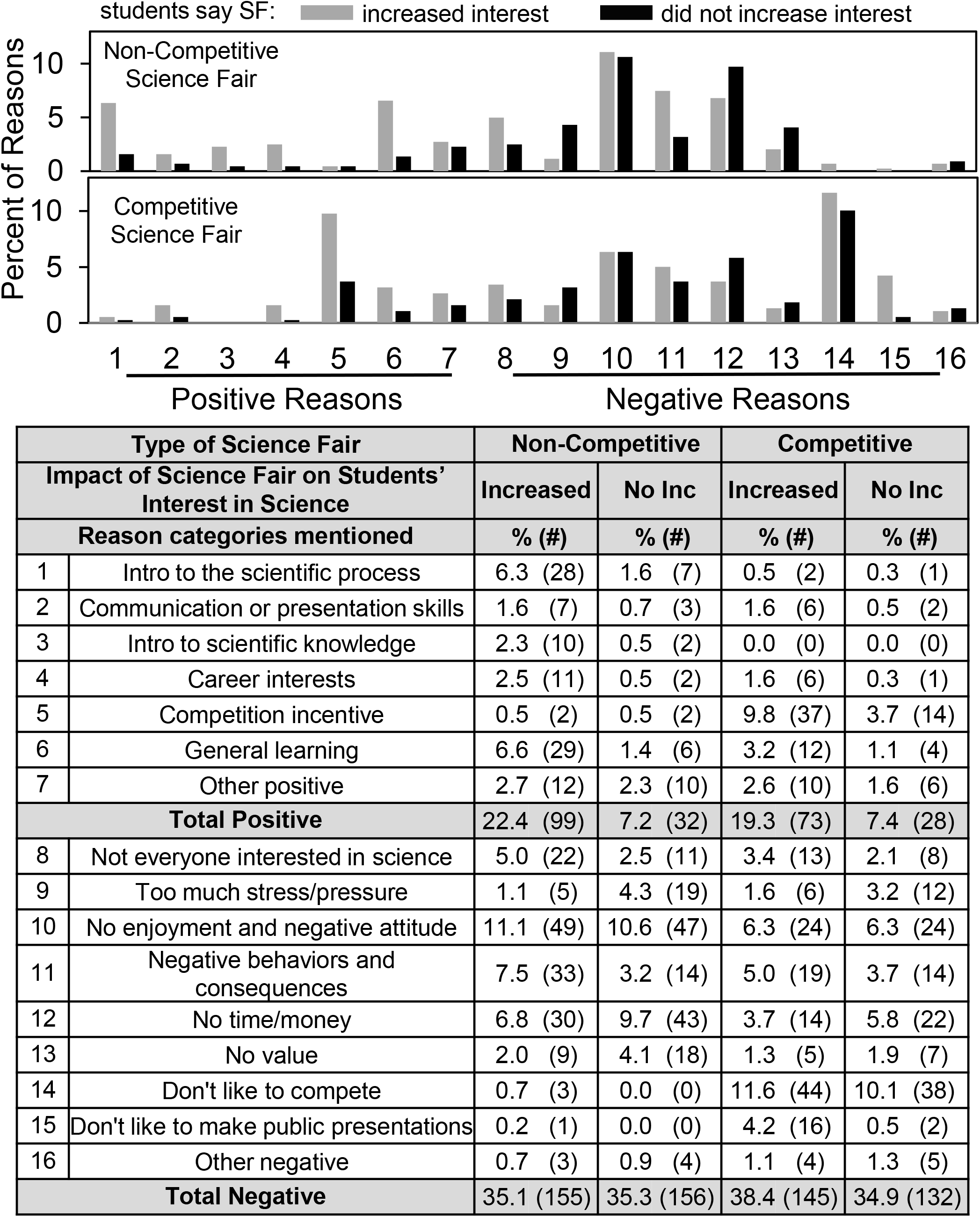
Distribution of student reasons positive and negative to require science fair depending on whether or not students say science fair increased their interest in science.

## Discussion

The goal of our ongoing research is to identify strengths and weaknesses of high school level science fair and improvements that can help science educators make science fair a more effective, inclusive and equitable learning experience. Previously, we reported findings regarding students’ high school science fair experiences based on a regional group of high school students who had competed in the Dallas Regional Science and Engineering Fair (DRSEF) and post high school students on STEM education tracks doing research at UT Southwestern Medical Center [10, 11]. In the current paper, we confirm and extend the previous findings. We added new questions to our anonymous and voluntary surveys to learn the extent to which students had an interest in science or engineering careers and if science fair participation increased their interest in science or engineering. And we surveyed a national group of high school students by incorporating our survey into the Scienteer (www.scienteer.com) online portal now used Texas and some other states for science fair registration, parental consent, and project management.

More than 300 students completed surveys during 2017 and 2018, representing about 0.5% of the students that participated in high school science fair via Scienteer. Student demographics and answers to questions regarding sources of help, types of help received, obstacles encountered, and ways of overcoming obstacles were very similar comparing the 2017 and 2018 responders. That more than 80% of the 2017/2018 national students wrote thoughtful answers to the open-ended text questions was an indication that the students took the surveys seriously. The finding that >95% of the positive student comments about science fair were given by the 20-25% of students who said that science fair should be required also provided validation of the survey responses.

A potential limitation of our study is the small size of the study population relative to the total number of students participating. Nevertheless, we observed many overlapping features of science fair experience between the low response rate/large data set (Scienteer national) compared to the high response rate/small data set (DRSEF regional) described previously [10, 11]. Articles on the internet, teachers, and parents were the main sources of help; time pressure and coming up with the idea were the main obstacles; more background research and perseverance were the main ways to overcome obstacles; fine-tuning the report and developing the idea were important types of help received. This similarity supports our previous conclusion that many features of science fair are common to students notwithstanding the diversity of science fair formats.

One major difference between the national and region groups concerned the requirement to participate in science fair, that is, 68% of the national students vs. 8% of the regional students. The finding that 68% of the national students were required to participate in science fair suggests that NSTA guidance about voluntary student participation [9] is widely ignored, at least from the students’ perception. We cannot tell if the students understand “required to do science fair” differently from their schools’ intentions, e.g., required to participate in science fair to get into an advanced class or to increase one’s grade is not exactly truly required but can be perceived that way.

Two other major differences between the national and regional groups – coaching for the interview and help from articles in books and magazines – may reflect the highly supportive practices to incentivize rather than require student science fair participation by the North Texas suburban school district where most of the regional students attended. Local school district support clearly can have an impact on some aspects of student science fair experience. Moreover, the same group of experiential differences along with receiving help from teachers and doing more background research to overcome obstacles was characteristic of students who said that science fair participation increased their interest in the sciences or engineering, and who reported more perseverance and less difficulty becoming motivated.

Compared to the regional students [11], the 2017/2018 national group of students showed some noteworthy differences in their open-ended text comments. For instance, they mentioned the positive value of science fair towards *general learning* (7.9% & 4.2%) as well as *intro to scientific process* (7.9% & 0.7%), whereas few of the regional students mentioned general learning as an outcome of science fair. Also, they mentioned as negative reasons too much stress/pressure (5.4% & 4.8%) and no value (6.1% & 3.2%), neither of which was emphasized by the regional DRSEF students. And the negative comment *don’t like to make public presentations* (0.2% & 4.8%) made by the 2017/2018 students might reflect directly the lower number of students who reported receiving coaching for the interview.

Overall, the findings with the 2017/2018 national group of students are consistent with idea that the students’ focus switches from competition to learning when thinking about competitive vs. non-competitive science fair. For instance, regarding competitive science fair, the top negative reasons given by students were *don’t like to compete* (22%) and *no enjoyment/overall negative attitude* (13%); the top positive reason was *competition incentive* (14%). By contrast, the most common negative reasons about non-competitive science fair were *no enjoyment/overall negative attitude* (22%) and *no time/money* (17%); the top positive reasons were *general learning* (7.9%) and *intro to scientific process* (7.9%).

The potential value of non-competitive science fair in which judges assess on a sliding scale student progress towards mastery of the different practices of science has been described by others, albeit not for high school students [21–24]. By emphasizing learning vs. competing, the non-competitive approach would be consistent with student motivation and goal orientation theory, i.e., mastery (competition with oneself with emphasis on understanding and improving skills and knowledge) vs. performance (competing with others with emphasis on demonstrating high ability and grades) [25–27].

Increasing student interest in science represents one of the most important potential positive outcomes of science fair. Previous research by others had shown that participating in science competitions helped to maintain high school student interest in pursuing science education and science careers albeit to a small extent, but those studies did not take into account whether or not students were required to do science fair [28–32]. Other research has analyzed student motivations and the benefits of participating in science fair, but here too the impact of requiring science fair participation was not taken into consideration [33–35].

Our data shows that being required to participate in science fair can have the practical consequence of decreasing the positive impact on students. We found that about 60% of the students surveyed said that participating in science fair increased their interest in the sciences or engineering. That number was significantly higher if the students had chosen to participate in science fair rather than been required to do so. Indeed, requiring science fair participation decreased the positive impact of science fair regardless whether or not the students said they were interested in a career in the sciences or engineering. In the worst case, ~10% of the students who said that they were not interested in a career and were required to do science fair engaged in research misconduct, i.e. copying their project from someone else or making up the data. None of the regional high school students in our previous study reporting making up their data [10], but few were required to participate in science fair as has been discussed. On the other hand, 24% (5 of 21) of students, all of whom were required to participate in the 2000 Bell Montreal regional science fair, were reported to make up their data [14]. Taken together, the foregoing findings emphasize that requiring students to participate in science fair can have a negative outcome. Perhaps an analogous situation occurs when professional scientists perceive the institutional environment are unfair and, as a result, are more likely to engage in research misconduct [36].

In conclusion, our results lend strong empirical support to NSTA guidance that participation in science competitions should be voluntary [9]. The challenge will be for school districts to find ways to incentivize an activity that requires so much time and effort. Our findings also suggest that offering students a noncompetitive science fair option could provide a way to promote the NSTA goal that science fair emphasis should be on the learning experience rather than the competition and would be an especially important option for students who do not want to compete. Finally, the availability of two kinds of science fairs -- competitive and non-competitive -- may help achieve the dual objectives of science education -- science for the scientists and engineers of the future and science for everyone [37]. Recently, we put forth these policy ideas in a commentary in NSTA Reports [12]. In future studies, we hope to gain further insights about student science fair experience though new survey questions that we have added regarding high school geographic location and student ethnicity.

## Acknowledgments

We are grateful to Russell Cowen and Rocky Slavin, managers of Scienteer Technologies, who incorporated the parental consent and science fair survey REDCap link into the Scienteer website and continue to provide ongoing oversight and management of survey access. Use of REDCap survey and data management tool was facilitated by the UTSW Department of Population and Data Sciences and Clinical and Translational Science Training Program. Karen Shepherd (Plano Independent School District) and Dr. Ann Batenburg (Southern Methodist University) helped with survey development. Dr. Shannon A. Scielzo (UT Southwestern) advised us about survey metrics.

## Supporting information

**S1 Data set.** Excel dataset showing all of the survey questions and answers for 2017 surveys.

**S2 Data set.** Excel dataset showing all of the survey questions and answers for 2018 surveys.

**S3 Data set.** Excel dataset showing the complete set of reason category assignments.

**S1 Survey**. Survey questions.

**S1 Fig.**
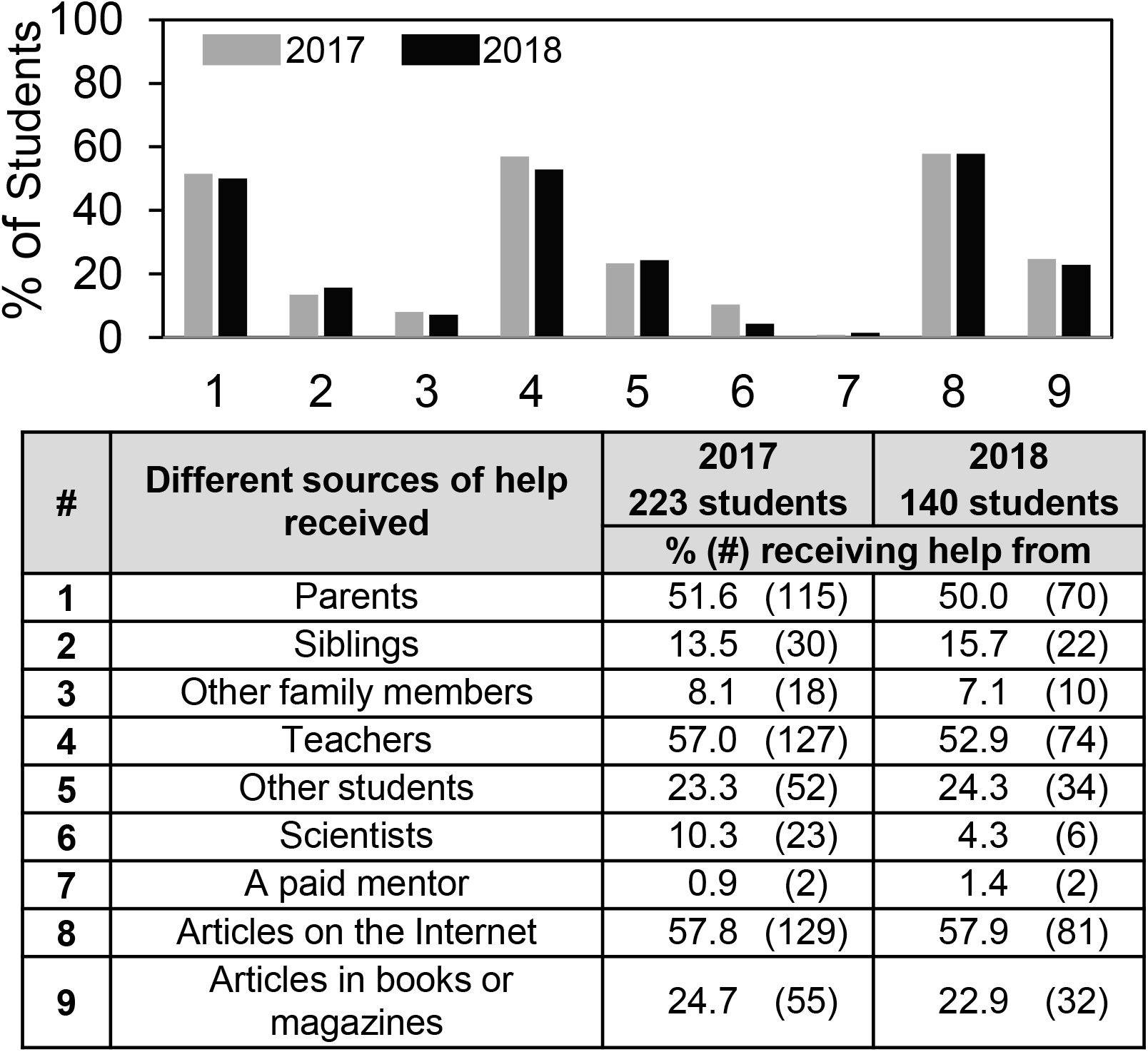
Frequency of student answers to the question “Who helped you with your science fair project?”

**S2 Fig.**
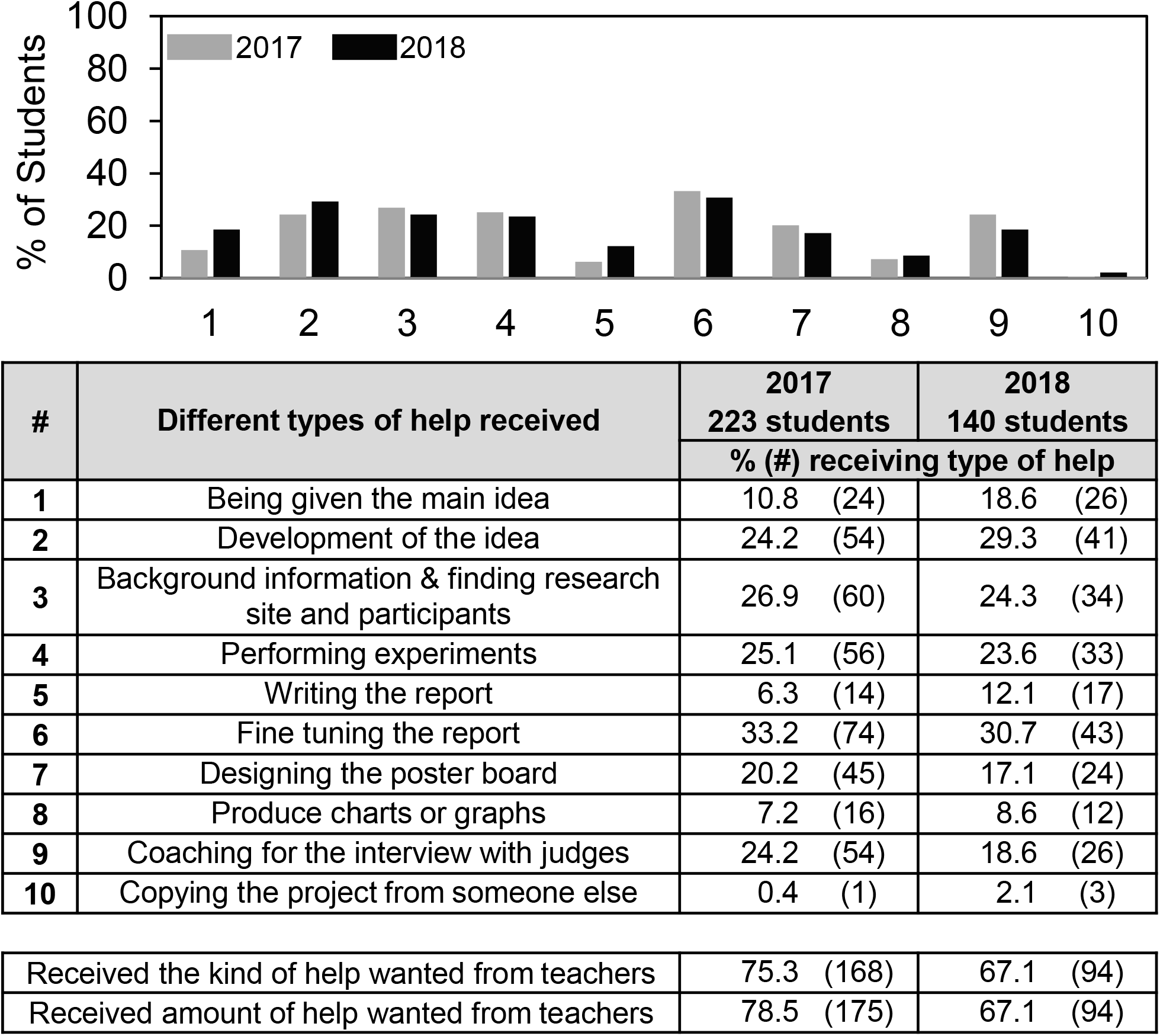
Frequency of student answers to the question “What kind of help did you receive doing science fair?”

**S3 Fig.**
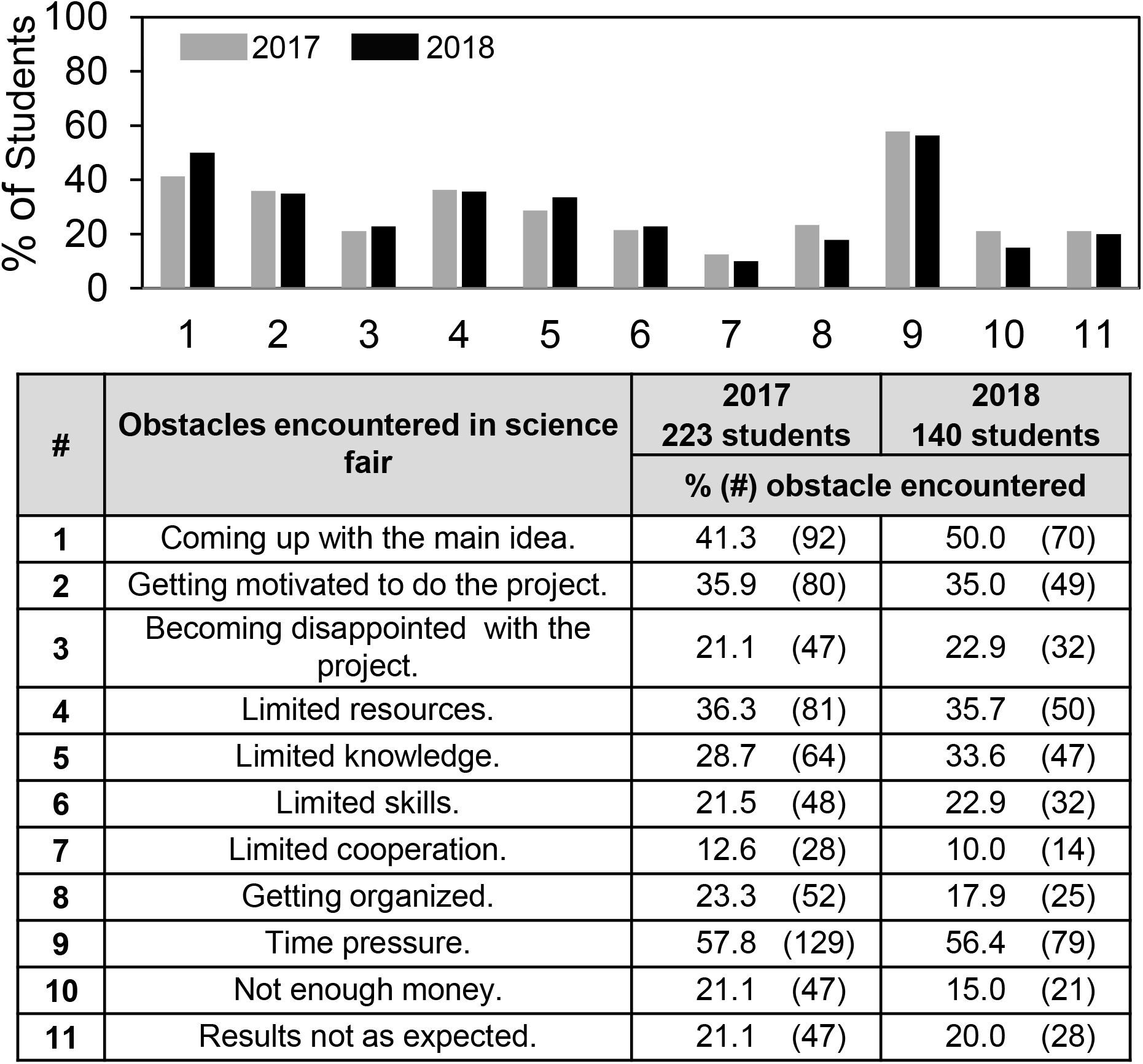
Frequency of student answers to the question “In your science fair project, what obstacles did you face?”

**S4 Fig.**
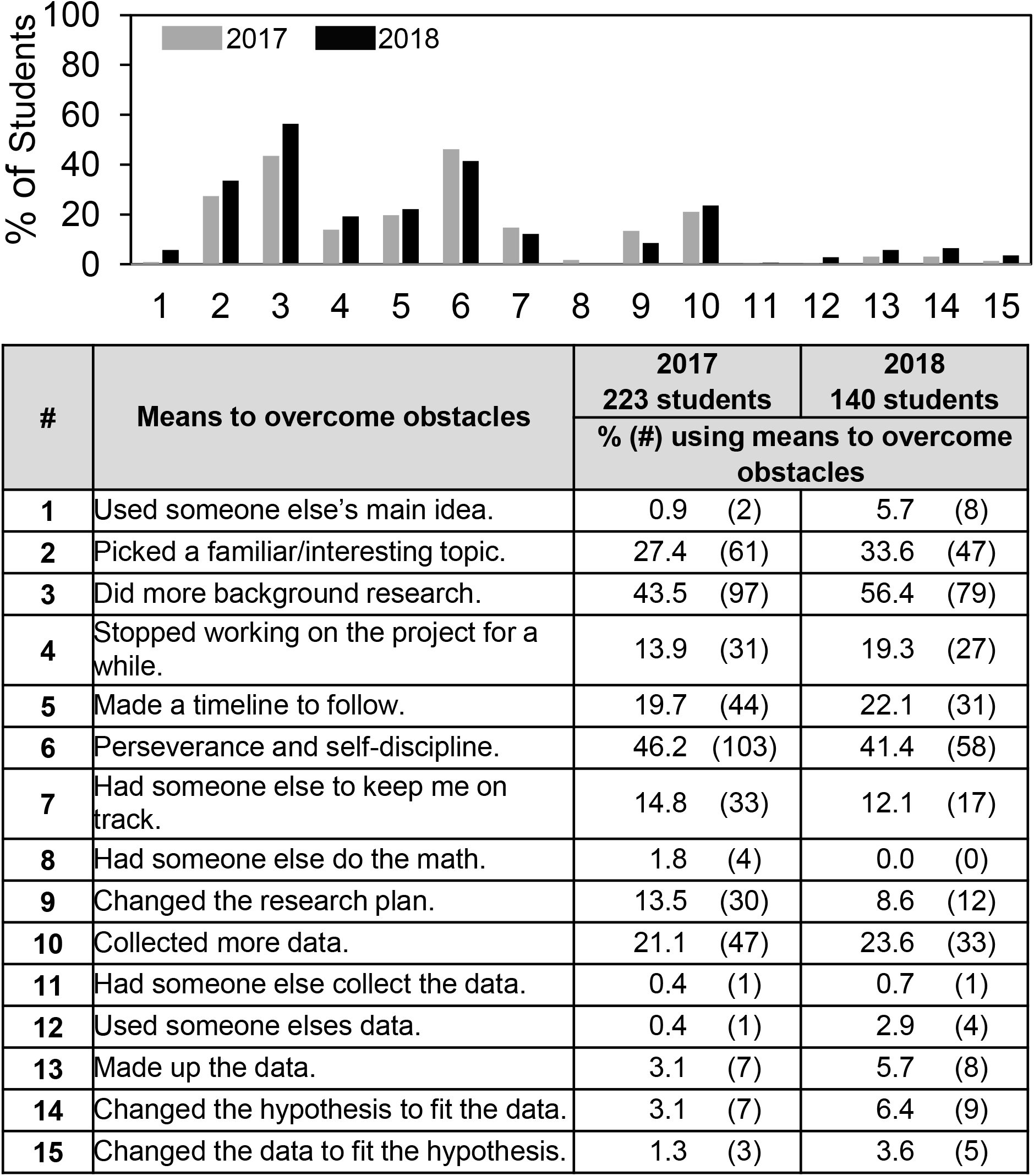
Frequency of student answers to the question “In your science fair project, how did you overcome obstacles?”

**S5 Fig.**
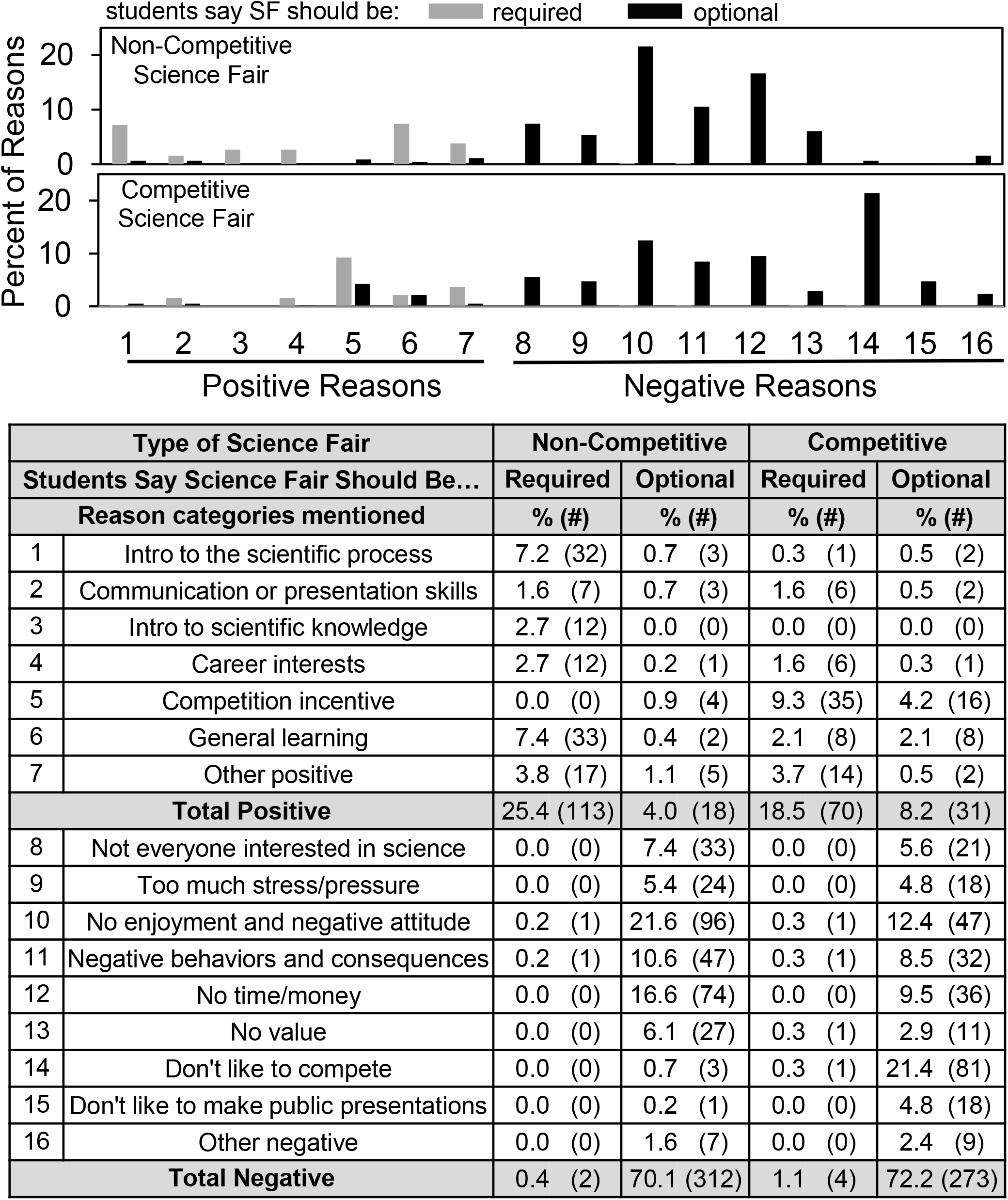
Distribution of reasons positive and negative to require science fair depending on whether students said science fair should be required or optional.

## Notes

#### Summary of Updates

Revised manuscript in response to PLOS ONE editorial and referee comments. Corresponds to the new version re-submitted to PLOS ONE.

